# Stacking multiple optimal transport policies to map functional connectomes

**DOI:** 10.1101/2023.02.06.527388

**Authors:** Javid Dadashkarimi, Matthew Rosenblatt, Amin Karbasi, Dustin Scheinost

## Abstract

Connectomics is a popular approach for understanding the brain with neuroimaging data. However, a connectome generated from one atlas is different in size, topology, and scale compared to a connectome generated from another. Consequently, connectomes generated from different atlases cannot be used in the same analysis. This limitation hinders efforts toward increasing sample size and demonstrating generalizability across datasets. Recently, we proposed Cross Atlas Remapping via Optimal Transport (CAROT) to find a spatial mapping between a pair of atlases based on a set of training data. The mapping transforms timeseries fMRI data parcellated with an atlas to form a connectome based on a different one. Crucially, CAROT does not need raw fMRI data and thus does not require re-processing, which can otherwise be time-consuming and expensive. The current CAROT implementation leverages information from several source atlases to create robust mappings for a target atlas. In this work, we extend CAROT to combine existing mappings between a source and target atlas for an arbitrary number of mappings. This extension (labeled Stacking CAROT) allows mappings between a pair of atlases to be created once and re-used with other pre-trained mappings to create new mappings as needed. Reconstructed connectomes from Stacking CAROT perform as well as those from CAROT in downstream analyses. Importantly, Stacking CAROT significantly reduces training time and storage requirements compared to CAROT. Overall, Stacking CAROT improves previous versions of CAROT.

## 1 Introduction

Functional magnetic resonance imaging (fMRI) data can be represented as a functional connectome, which is a matrix describing the connectivity between any pair of brain regions. It is created by parcellating the brain into distinct regions using an atlas and estimating their connectivity (*e.g*., by calculating correlations between timeseries data for each pair of regions). Different atlases divide the brain into different regions of varying size and topology. As such, connectomes created from different atlases are not directly comparable. Since numerous atlases exist [2], the inability to directly compare connectomes hinders cross-study interpretation, replication, generalization, and data-pooling.

One solution to this problem is to create a mapping between connectomes from different atlases. With such a mapping, connectomes from one atlas can be transformed into connectomes from another without the need to reprocess “raw” fMRI data. We previously developed Cross Atlas Remapping via Optimal Transport (CAROT) to create these mappings [8, 7, 9]. CAROT uses optimal transport, or the mathematics of converting a probability distribution from one set to another, to find a spatial mapping between a pair of atlases based on a set of training data. The mapping transforms timeseries fMRI data parcellated with the first atlas (source atlas) to form a connectome based on the second atlas (target atlas). Crucially, CAROT does not need raw fMRI data and thus does not require re-processing, which can otherwise be time-consuming and expensive. Not only do original and CAROT-reconstructed connectomes have similar features, but they also perform similarly in downstream analyses [8, 7, 9].

The current CAROT implementation leverages information from several source atlases to create robust mappings for a target atlas. A key limitation of this implementation is that data from all atlas pairs are needed during the initial training of the connectome mappings. In other words, if a new atlas is introduced or an atlas is removed, the mappings must be retrained. In this work, we extend CAROT to combine existing mappings between a source and target atlas for an arbitrary number of mappings. This extension allows mappings between a pair of atlases to be created once and re-used with other pre-trained mappings to create new mappings as needed. Using six different atlases, we show that our approach maintains the high quality of reconstructed connectomes (equal to that of the current CAROT implementation) while decreasing training time and storage requirements.

## 2 Background

### 2.0.1 Optimal transport

The optimal transport problem solves how to transport resources from one location α to another *β* while minimizing the cost *C* to do so [25, 15, 18, 13]. Using a probabilistic approach in which the amount of mass located at *x_i_* potentially dispatches to several points in the target distribution [17], admissible solutions are defined by a coupling matrix 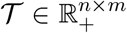 indicating the amount of mass being transferred from location *x_i_* to *y_j_* by 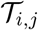:

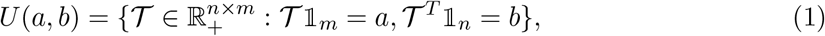

for vectors of all 1 shown with 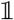. An optimum solution is obtained by solving the following problem for a given “ground metric” matrix 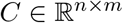 [21]:

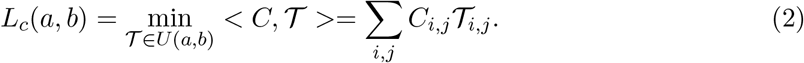

which is a linear problem and is not guaranteed to have a unique solution [19]. Though, it always has an optimal solution (see proof in [4, 3]). Unlike the KL divergence or other approaches, optimal transport provides a well-defined distance metric when the support of the distributions is different.

### 2.0.2 Single-source CAROT

In its simplest form, CAROT transforms the timeseries data from one atlas (labeled the source atlas) into timeseries data from an unavailable atlas (labeled the target atlas). Next, the corresponding functional connectomes are estimated using standard approaches (e.g., full or partial correlation). Formally, it is assumed that we have training timeseries data consisting of *T* timepoints from the same individuals but from two different atlases (atlasP_n_ with *n* regions and atlasP_m_ with *m* regions). Additionally, let 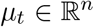 and 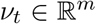 be the vectorized brain activity at single timepoint *t* based on atlasesP_n_ andP_m_, respectively. For a fixed cost matrix 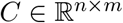, which measures the pairwise distance between regions inP_m_ andP_n_, this approach aims to find a mapping 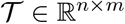 that minimizes transportation cost between *μ_t_* and *ν_t_*:

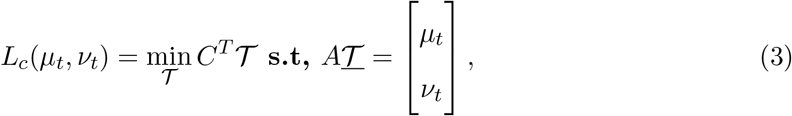

in which 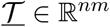 is vectorized version of 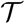 such that the *i* + *n*(*j* – 1)’s element of 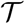 is equal to 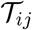 and *A* is defined as:

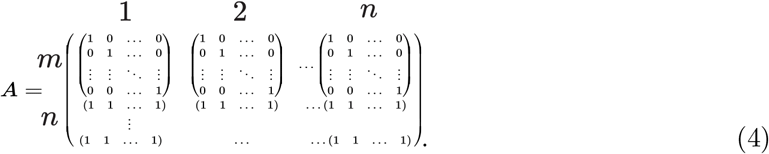

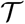 represents the optimal way of transforming the brain activity data from *n* regions into *m* regions. Thus, by applying 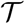 to every timepoint from the timeseries data of the source atlas, the timeseries data of the target atlas and corresponding connectomes can be estimated. The cost matrix *C* was based on the similarity of pairs of timeseries from the different atlases:

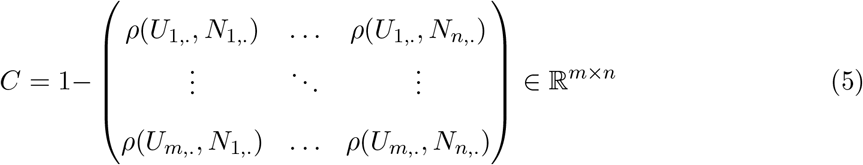

where *U_x_* and *N_x_* are timeseries fromP_m_ andP_n_ and *ρ*(*U_x_, N_y_*) is correlation between them.

### 2.0.3 Multi-source CAROT

When multiple source atlases exist (which is common), the above solution can be generalized to leverage information from each source atlas. In general, assume we have paired timeseries, from the same person, but from *k* different source atlases with a total of *n_s_* regions (where *n_s_* = *n*_1_ + *n*_2_ +.. + *n_k_* from source atlasP_n_1__ with *n*_1_ regions,P_n_2__ with *n*_2_ regions,.., P_nk_ with *n_k_* regions) and a target atlasP_m_ with *m* regions. Let us define 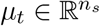 and 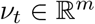 to be the distribution of brain activity at single time point *t* based on atlasesP_s_ andP_m_:

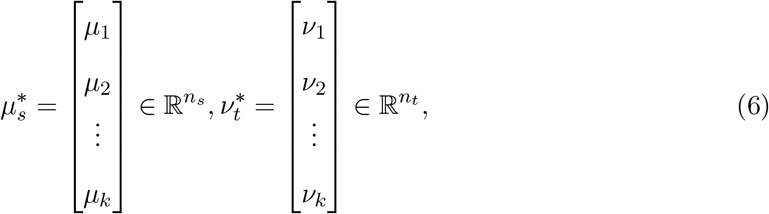

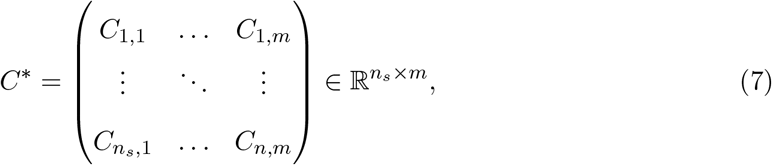

and *C_i,j_* is based the similarity of pairs of timeseries from nodes *i* and *j* from different atlases. Next, we want to minimize distance between 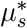 and *v_t_* as:

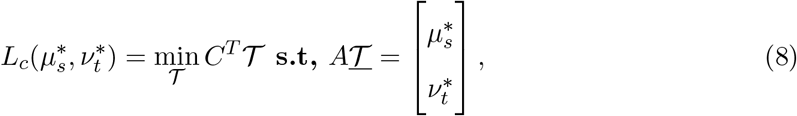

## 3 Stacking CAROT

Although Multi-source CAROT successfully leverages information from multiple source atlases to improve reconstruction, it has significant training and storage requirements. Here, we suggest a more scalable architecture, Stacking CAROT, to maximize flexibility and usability.

### 3.0.1 Limitations of multi-source CAROT

One limitation of training a large policy is the amount of training time. Training a large matrix takes longer than training a smaller one, given that we should calculate it each time we want to transfer mass. Furthermore, the matrix configuration could differ depending on the number of available source atlases. For example, for a set of *n* – 1 available atlases, the total possibilities to configure this problem equals:

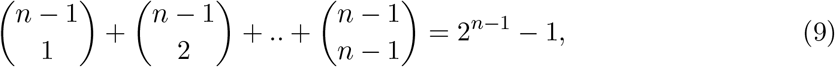

which equals the total number of subsets of a set with size=*n* – 1 except for the empty set.

### 3.0.2 Stacking CAROT

Stacking CAROT concatenates previously trained transportation plans to create a new mapping from *n* inputs to one output. Thus, in contrast to Multi-source CAROT, Stacking CAROT only needs to train *n* policies for the same number of target atlases as it is flexible to any source configurations. Let’s define training steps for each pair of source and target atlas:

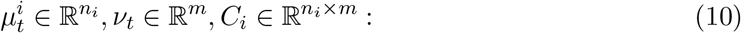

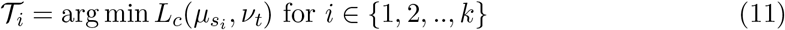

and in testing we want to estimate 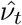 as follows:

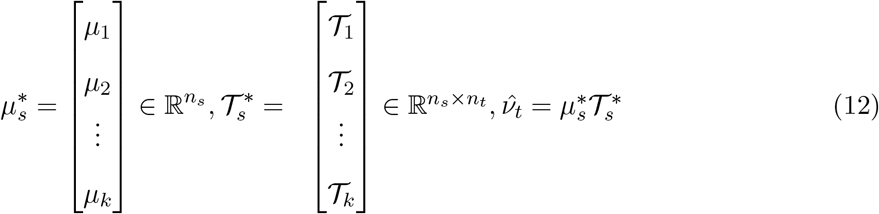

## 4 Implementation

Solving the large linear program in Equation 8 is computationally hard [10]. As such, we used entropy regularization, which gives an approximation solution with complexity of 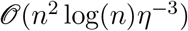 for 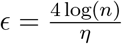 [19], and instead solve the following:

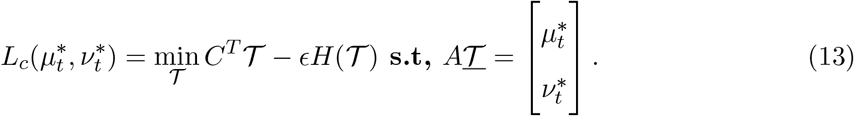

Specifically, we use the Sinkhorn algorithm—an iterative solution for Equation 13 [1]—to find 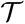, as implemented in the Python Optimal Transport toolbox [12].

## 5 Results

### 5.0.1 Datasets

To evaluate our approach, we used data from the Human Connectome Project (HCP) [26], starting with the minimally preprocessed data [14]. First, data with a maximum frame-to-frame displacement of 0.15 mm or greater were excluded, resulting in a sample of 876 restingstate scans. Analyses were restricted only to the LR phase encoding, which consisted of 1200 individual time points. Further preprocessing steps were performed using BioImage Suite [16]. These included regressing 24 motion parameters, regressing the mean white matter, CSF, and grey matter timeseries, removing the linear trend, and low-pass filtering. The Shen (268 nodes) [24], Schaefer (400 nodes) [22], Craddock (200 nodes) [6], Brainnetome (246 nodes) [11], Power (264 nodes) [20], and Dosenbach (160 nodes) [5] atlases were applied to the preprocessed data to create mean timeseries for each node. Connectomes were generated by calculating the Pearson’s correlation between each pair of these mean timeseries and then tasking the Fisher transform of these correlations.

### 5.0.2 Similarity between reconstructed and original connectomes

To validate Stacking CAROT, we assess the similarity of connectomes reconstructed using Stacking CAROT and the original connectomes generated directly from the raw data. First, we partitioned our sample into 80% training data to estimate the optimal mapping 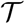 between atlases and 20% testing data to evaluate the reconstructed connectome quality. In the training data, we estimated 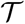 using all 1200 time points and 700 participants for each source-target atlas pairs. Next, we applied the estimated 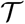 to reconstruct the target atlases in the testing data. Finally, the reconstructed connectomes were compared to the “gold-standard” connectomes (*i.e*., connectomes generated directly from an atlas) using correlation.

We compared these similarities from Stacking CAROT to the similarities from Multi-source CAROT. As in our prior work, multi-source CAROT was trained with the same 80/20 split and all available source atlases to a single target atlas. We hypothesize that Stacking CAROT will produce connectomes significantly correlated with the “gold-standard” connectomes and that no difference between Stacking CAROT and Multi-source CAROT would exist.

As shown in Figure 2, we observed significant correlations between the reconstructed connectomes and their original counterparts for both Stacking CAROT and Multi-source CAROT (all *ρ*′*s* > 0.50 and *p′s* < 0.05).

### 5.0.3 Impact on the number of atlases

We investigated the impact of using a smaller number of source atlases by only including *k* random source atlases when creating connectomes for the target atlas. This process was repeated with 100 iterations over a range of *k* =2–6. As shown in Figure 3, strong correlations (e.g., *ρ* > 0.6) can be observed with as little as two or three source atlases. No differences between Multi-source CAROT and Stacking CAROT were observed.

**Figure 1:**
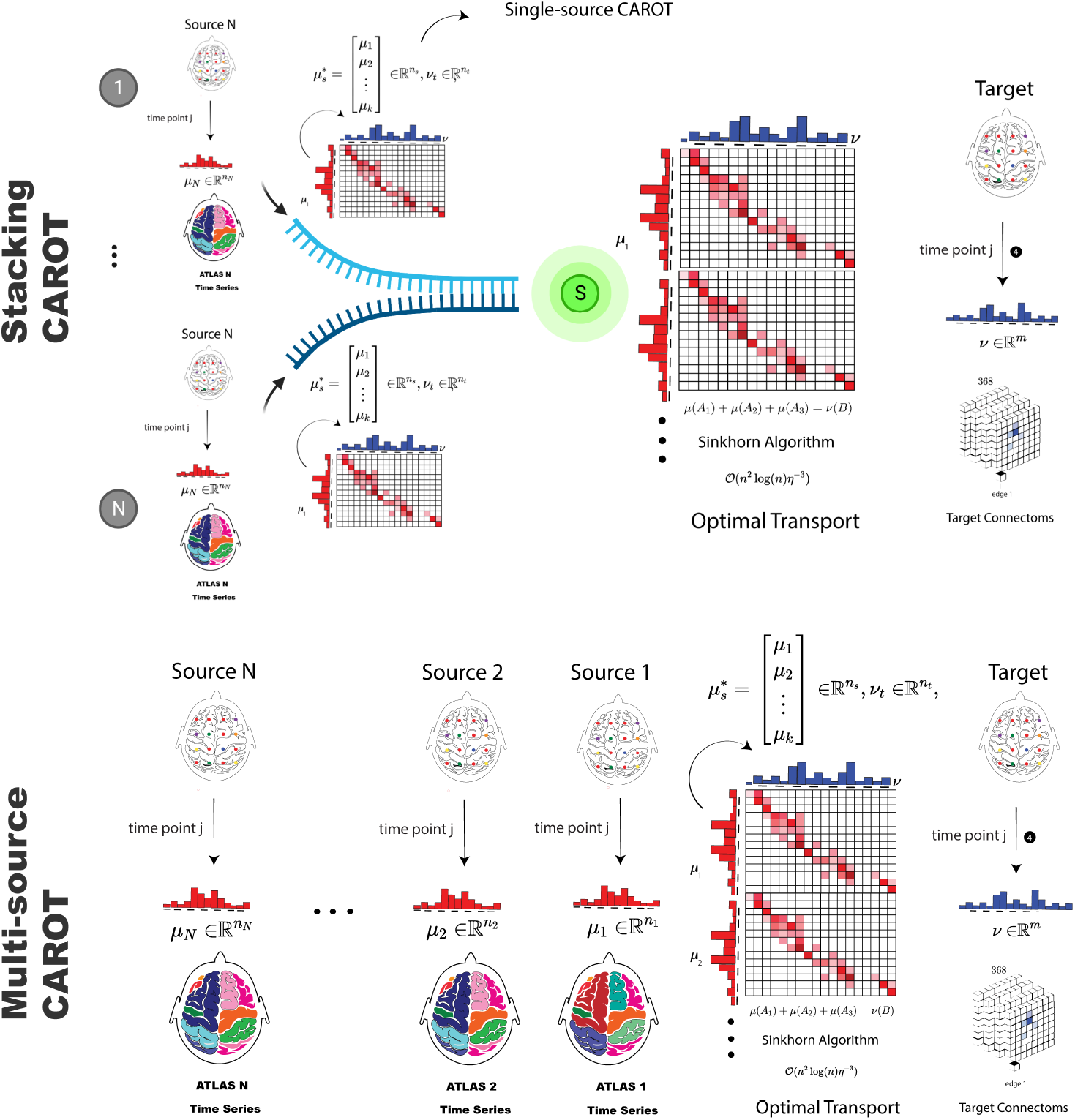
Our Stacking CAROT approach combines pre-trained mappings between atlas. Compared to the current CAROT implementation (*e.g*., Multi-Source CAROT), combining pre-trained models reduces training time and storage requirements.

**Figure 2:**
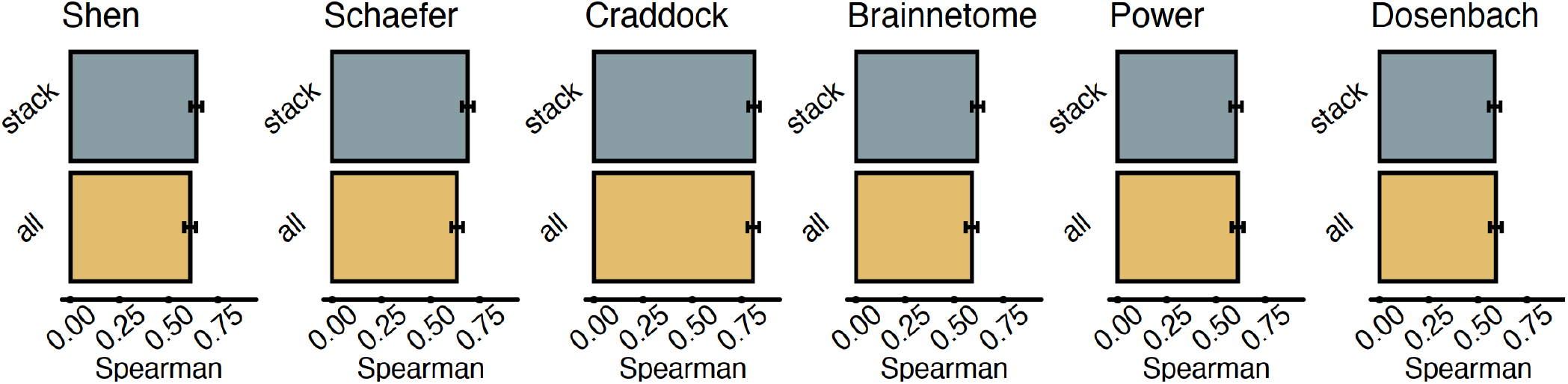
The Spearman’s rank correlation between the reconstructed connectomes and connectomes generated directly with the target atlases are shown Multi-source CAROT and (*bottom*) Stacking CAROT. No differences between the two approaches were observed. Error bars are generated from 100 iterations of randomly splitting the data into training and testing data.

**Figure 3:**
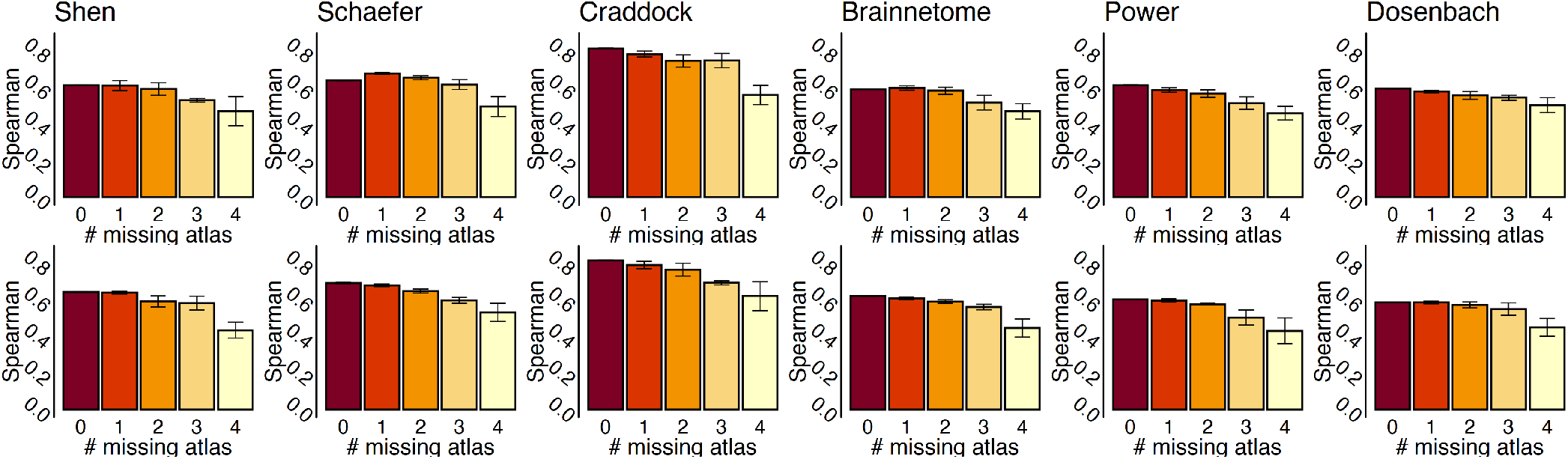
Bar plots exhibit the correlation between estimated connectomes and original connectomes based on *n* – *k* samplings of available atlases (*i.e*., *n* indicates the number of all available atlases to be transported) for each target atlas for (*top*) Multi-source CAROT and (*bottom*) Stacking CAROT. Both approaches can show strong correlations with less than the maximum number of source atlases. No differences between the two approaches were observed.

### 5.0.4 IQ prediction

Furthermore, we show that reconstructed connectomes can predict fluid intelligence using connectome-based predictive modeling (CPM) [23]. We partitioned the HCP dataset into three groupings: *g*_1_, which consisted of 25% of the participants; *g*_2_, which consisted of 50% of the participants; and, *g*_3_, which consisted of the final 25% of the participants. In *g*_1_, 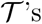 for each algorithm were estimated as above. We then applied 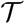 on *g*_2_ and *g*_3_ to reconstruct connectomes. Finally, for each set of connectomes, we trained a CPM model of fluid intelligence using *g*_2_ and tested this model in *g*_3_. Spearman correlation between observed and predicted values was used to evaluate prediction performance. This procedure was repeated with 100 random splits of the data into the three groups. In all cases, connectomes reconstructed using all of the source atlases performed as well in prediction as the original connectomes (Figure 4).

**Figure 4:**
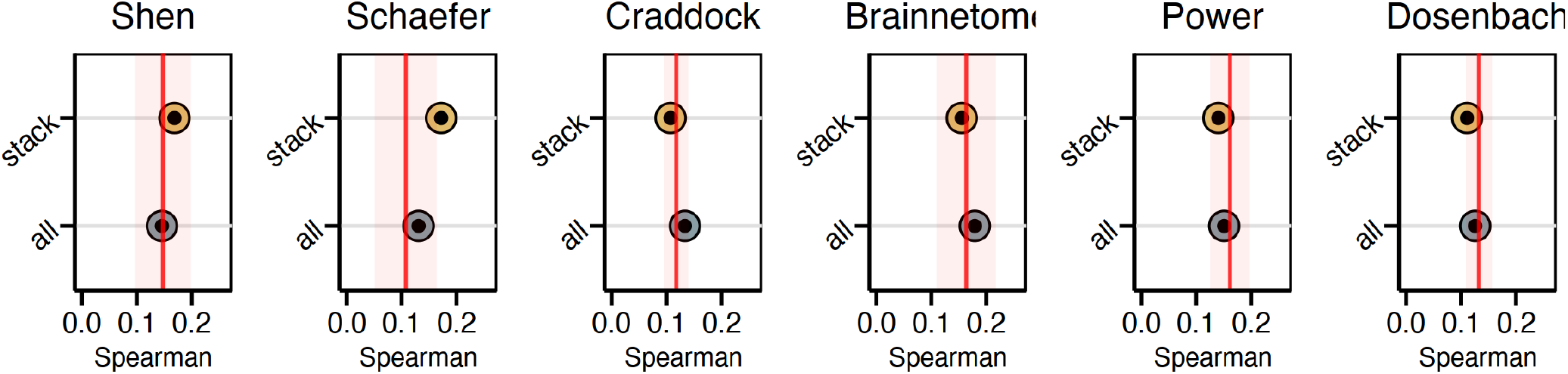
Multi-source and Stacking CAORT equally predicted IQ compared to each other as well as the original connnectome (red line). Size of circle represents the variance of prediction performance of 100 iteration of 10-fold cross-validation.

### 5.0.5 Stacking CAROT reduces training time and storage requirements

The main benefit of Stacking CAROT is the reduction of training time. For example, training Mutli-source CAROT with 5 atlases takes 100 minutes due to the large cost matrix *C*\*. In contrast, training CAROT with a single source takes 2 minutes. Thus, the total training time for Stacking CAROT with five atlases is 5 * 2 = 10 minutes. Nevertheless, the reduction in training time with Stacking CAROT is underappreciated in this example. If one had four source atlases (instead of five), a new Multi-source CAROT model would need to trained, requiring 40 minutes to train. However, as Stacking CAROT simply combines already trained models, there is no extra training time. Importantly, as applying the Multi-source or Stacking CAROT model is a frame-by-frame multiplication of the same dimension, the run time for applying a model to reconstruct a connectome is the same for both.

Similarly, Stacking CAROT is more efficient in terms of storage requirements. A single source model is 1.5MB. Thus, as models are combined as needed in run time, Stacking CAROT needs 30 * 1.5MB = 45MB to store all possible single source models for the six atlases we tested. In contrast, Multi-source CAROT needs greater than 3GB of storage.

## 6 Discussion and conclusions

Here, we significantly improve upon CAROT by implementing a novel stacking approach. Stacking CAROT combines previously estimated mappings between a single source and target atlas, allowing these previously estimated mappings to be reused and combined when multiple atlases are available. Reconstructed connectomes from Stacking CAROT perform as well as those from Multi-source CAROT in downstream analyses. Importantly, Stacking CAROT significantly reduces training time and storage requirements compared to Multi-source CAROT. Future work includes generalizing our framework to other functional timeseries data—*e.g*., electroencephalography (EEG) and functional near-infrared spectroscopy (fNIRS). Overall, Stacking CAROT is a promising extension to CAROT aimed to increase the generalization of connectome-based results across different atlases.

## Acknowledgment

This research study was conducted retrospectively using human subject data made available in open access by the Human Connectome Project. Approval was granted by the local IRB. Yale Human Research Protection Program (HIC #2000023326) on May 3, 2018. Data were provided by the Human Connectome Project, WU-Minn Consortium (Principal Investigators: David Van Essen and Kamil Ugurbil; U54 MH091657) and funded by the 16 NIH Institutes and Centers that support the NIH Blueprint for Neuroscience Research; and by the McDonnell Center for Systems Neuroscience at Washington University. Amin Karbasi is partially supported by NSF (IIS-1845032), ONR (N00014-19-1-2406), and Tata.

